# Drug connectivity mapping and functional analysis reveals therapeutic small molecules that differentially modulate myelination

**DOI:** 10.1101/2020.09.05.284372

**Authors:** F Pieropan, AD Rivera, G Williams, F Calzolari, AM Butt, K Azim

## Abstract

Oligodendrocytes are the myelin forming cells of the central nervous system (CNS) and are generated from oligodendrocyte progenitor cells (OPCs). Disruption or loss of oligodendrocytes and myelin has devastating effects on CNS function and integrity, which occurs in diverse neurological disorders, including Multiple Sclerosis (MS), Alzheimer’s disease (AD) and neuropsychiatric disorders. Hence, there is a need to develop new therapies that promote oligodendrocyte regeneration and myelin repair. A promising approach is drug repurposing, but most agents have potentially contrasting biological actions depending on the cellular context and their dose-dependent effects on intracellular regulatory pathways. Here, we have used a combined drug connectivity systems biology and neurobiological approach to identify compounds that exert positive and negative effects on oligodendroglia, depending on concentration. Notably, LY294002, a potent inhibitor of PI3K/Akt signalling, was the most highly ranked small molecule for both pro- and anti-oligodendroglial effects. We validated these *in silico* findings in multiple *in vivo* and *ex vivo* neurobiological models and demonstrate that low and high doses of LY294002 have a profoundly bipartite effect on the generation of OPCs and their differentiation into myelinating oligodendrocytes. Finally, we employed transcriptional profiling and signalling pathway activity assays to determine cell-specific mechanisms of action of LY294002 on oligodendrocytes and resolve optimal *in vivo* conditions required to promote myelin repair. These results demonstrate the power of multifactorial neurobiological and *in silico* strategies in determining the therapeutic potential of small molecules in neurodegenerative disorders.

**One-sentence summary:** Drug discovery and CNS myelination

## Introduction

In the CNS, myelin is produced by oligodendrocytes that are generated from oligodendrocyte progenitor cells (OPCs) throughout life (*1*). During postnatal development, in addition to generating neural progenitors (NP), neural stem cells (NSCs) of the subventricular zone (SVZ) also pass through a number of distinct differentiation stages to generate OPCs, which migrate throughout the forebrain and differentiate into myelinating oligodendrocytes, in response to intrinsic and extracellular cues (*2, 3*). In the adult brain, a significant population of endogenous OPCs persist throughout the brain and have the function of life-long generation of oligodendrocytes, which is essential for myelination of new connections in learning and myelin repair in pathology (*4, 5*). In addition, the adult SVZ remains an important source of new OPCs to replenish endogenous populations, in particular following pathological demyelination (*3, 6*). However, long-term repair ultimately fails due to a decline in oligodendrocyte regeneration from both the SVZ (*3, 7*) and endogenous OPCs (*8, 9*), which severely impairs repair in numerous neuropathologies, including Multiple Sclerosis (MS) and Alzheimer’s disease (AD) (*9, 10*). Hence, there is a need for new therapies that promote OPC regeneration and repair.

Connectivity mapping has been used in multiple clinical areas to connect biology and drug discovery by exploiting transcriptional similarities across treatment conditions and cell states (*11*). This strategy is a promising and direct approach to regulate neural cells and identify small molecules and transcriptional networks that have the potential to promote regeneration and repair in the CNS (*3, 12–14*). However, agents identified by these new therapeutic approaches have potentially divergent biological actions depending on their dose- and time-dependent effects on intracellular regulatory pathways. Thus, determining the cell-specific effects and precise mechanisms of action of small molecules on neural cells is essential for developing therapeutic strategies to promote CNS repair. In the present study, we have identified small molecules with the potential to regulate oligodendrocyte regeneration by utilizing a new comprehensive reference catalogue, LINCS (Library of Integrated Network-based Cellular Signatures), which hosts the gene expression phenotypes triggered by small molecules assayed at different concentrations across diverse cellular systems (https://clue.io). We identified a wide range of small molecules that targeted multiple regulatory pathways with the potential to both positively and negatively regulate oligodendrocyte differentiation. Significantly, the highest ranking small molecule, LY294002, was predicted to have both pro- and anti-oligodendroglial actions. LY294002 is a potent inhibitor of the PI3K/Akt signalling pathway, which is considered essential for oligodendrocyte differentiation and myelination (*15, 16*). We validated the pharmacogenomic findings in multiple *in vivo* and *ex vivo* neurobiological models and demonstrate for the first time that LY294002 has a striking dose-dependent effect on oligodendrocytes, being severely destructive at high doses, but greatly stimulating oligodendrocyte generation and myelination at low doses. Furthermore, using whole genome transcriptomics and biochemical assays, we determined the cell-specific differential mechanisms of action of low and high LY294002 in oligodendrocytes. This study identifies striking contrasting effects of small molecules on neural cells depending on their dose-dependent actions on intracellular regulatory pathways, which is critical for the development of novel therapeutic strategies using small molecules to promote CNS repair.

## Results

### Pharmacogenomic screening for small molecules predicted to regulate myelination

First, using datasets generated by the authors to profile postnatal and adult NSC and OL lineage cells (*3, 13, 17–20*), we curated essential landmark genes that can be defined as pro- or anti-oligodendrocyte differentiation (Fig. 1A). Next, we interrogated these genes in LINCS (https://clue.io) (*11*), to identify small molecules that shift the transcriptome of IPSC-derived NSCs (iNSCs) to that of myelinating oligodendrocytes (Fig. 1B). Then, we analysed the small molecule target-genes (TGs) whose expression is significantly modulated in iNSC to either positively or negatively regulate oligodendrocyte differentiation (Fig. 1C, D). Many of these TG networks have recognised functions in oligodendrocytes, such as the pro-oligodendroglial effects of inhibiting GSK3β (*21*) and anti-oligodendroglial effects of inhibiting mTOR (*22*). In contrast, the pro-oligodendroglial effect of HDAC inhibition appears counter-intuitive at first, since HDAC activity is essential for oligodendrocyte lineage progression (*23*), but transient HDAC inhibition can be neuroprotective and promote OPC plasticity (*24, 25*). Significantly, inhibition of a number of pathways is predicted to have both pro- and anti-oligodendrocyte functions, most notably PI3K/Akt/mTOR inhibition (Fig. 1D). Consistent with this, the highest ranking pro- and anti-oligodendroglial small molecule was LY294002, a potent inhibitor of PI3K/Akt signalling, with broad kinase activity depending on concentration and cellular context (*26*). LY294002 exerts opposing transcriptional effects on iNSCs at low concentrations (2 μM, here termed L-LY29) and high concentrations (10 μM, here termed H-LY29) (Fig. 1E). The second highest-ranking anti-oligodendroglial small molecule was the specific Akt inhibitor Triciribine (TCN, Fig. 1E), and comparison of the target gene (TG) pathways and biological processes altered by the non-specific PI3K/Akt inhibitor LY294002 compared to the specific Akt inhibitor TCN provided insight into the potential mechanisms that determine the opposing effects of L- and H-LY29 on oligodendrocytes and myelination, with the “FAK-PI3K-mTOR pathway” being most prominent (Fig. S1). In support of this, meta-analysis of the LINCS-derived TGs closely associated H-LY29 with TCN (Fig. 1F), consistent with the potential anti-oligodendroglial actions of H-LY29 being due to inhibition of PI3K/Akt/mTOR signalling, which is critical for OPC differentiation and myelination (*27*). In contrast, the pro-oligodendroglial actions of L-LY29 are most closely linked to the effects exerted by GSK3β inhibitors and metformin (Fig. 1F), both of which have been shown to rejuvenate OPC regeneration and promote remyelination (*3, 9, 21*), together with corticosteroids and the flavonoid epicatechin. Interestingly, L-LY29 is also closely associated with perturbation of HDACs (Fig. 1F), which has broad spectrum epigenetic effects that regulate OPC plasticity (*25*). These analyses identify potential mechanisms that determine predicted opposing effects of H- and L-LY29 on oligodendrocytes and myelination

**Fig. 1:**
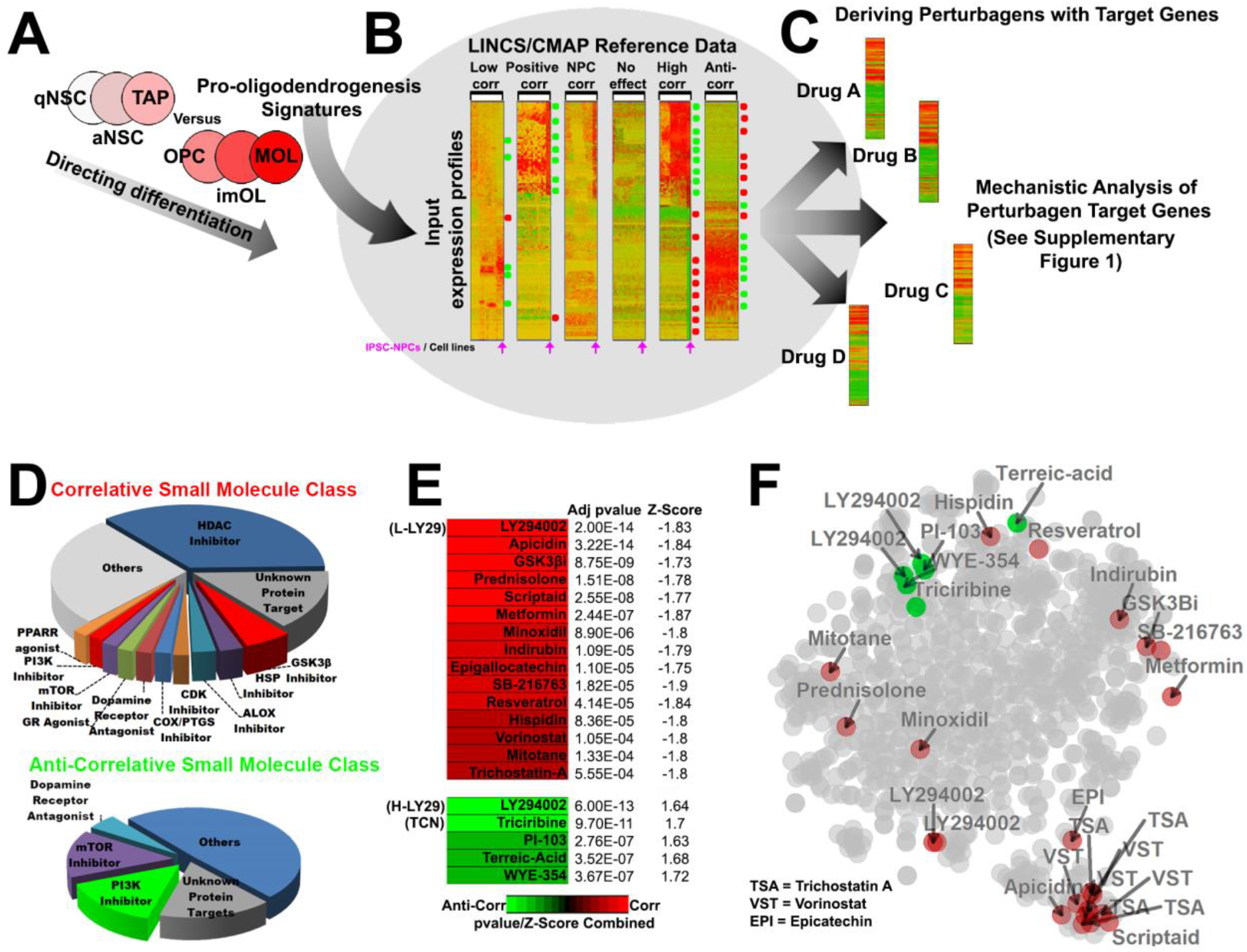
Querying the LINCS database for small molecules modulating oligodendroglia and insights into their cellular mechanisms of action. (**A,B**) Transcriptional signatures of early and late OL lineage stages are used to query the LINCS database, which contains drug-induced expression profiles for over 20,000 small molecules assayed in over 90 cells lines. To focus on the most OL-relevant data, only LINCS datasets comprising IPSCs-NSCs were queried. (**C**) Matching small molecule target genes (TGs) are extracted for subsequent mechanistic investigations. (**D**) The broader mechanisms of action/target proteins of the highest ranked small molecules. (**E**) Heatmap output of the top ranking small molecules predicted to enhance (red) or inhibit (green) myelination, sorted by their adjusted (adj) pvalues, coloured by their combined pvalue/z-score (correlation in input profiles with the profiles induced by IPSCs-NSCs within the database). Small molecules tested are abbreviated in brackets. (**F**) tSNE plot illustrating the distance/similarities in TGs induced by top ranking oligodendroglial perturbing small molecules. Note the distance between LY-294002 TGs (LY-29) for the higher (green) and lower (red) concentrations.

### Bipartite concentration-dependent effects of LY294002 on oligodendrocyte development *in vivo*

The pharmacogenomic analysis identified LY294002 as a potentially potent modulator of oligodendrogenesis and predicted differential effects of low and high concentrations. We tested this *in vivo* by direct injection of agents into the cerebrospinal fluid (CSF) of anesthetised mice for three days, commencing at postnatal day (P8) and analysing brains at P11, as described previously (*21*). These ages correspond to the main period of oligodendrocyte differentiation and myelination in the corpus callosum and dorsal cortex, after which the numbers of OPCs decline sharply by half with a concomitant doubling in newly formed myelinating oligodendrocytes (MYOLs) (*21, 28*). First, we determined the *in vivo* concentration-dependent effects of LY294002 and TCN (Fig. S2); TCN was selected because it is a selective small molecule inhibitor of Akt, but does not inhibit PI3K, the direct upstream activator of Akt (*29, 30*), whereas LY294002 is a potent inhibitor of PI3K, the upstream activator of Akt, but also has broad kinase activity (*26*). As predicted from the LINCS analysis, low doses of LY2940002 (≤2 μM) and high doses of LY294002 (≥10 μM) had contrasting pro- and anti-oligodendroglial effects in the corpus callosum, whereas TCN was anti-oligodendroglial at all concentrations tested (Fig. S2). Based on these doseresponse experiments, we performed a detailed *in vivo* analysis of the effects of 2 μM LY294002 (L-LY29) and 20 μM LY294002 (H-LY29), compared with 1.3 μM TCN on OPC, MYOL and myelination (Fig. 2). Immunolabelling for PDGFRα and the cell proliferation marker PCNA demonstrate that L-LY29 increased the number of OPCs in cell cycle and doubled their number overall, in both the corpus callosum and dorsal cortex (Fig. 2A, B, E, F). In contrast, H-LY29 significantly reduced OPC numbers and proliferation (Fig. 2C, E, F), whereas TCN did not decrease OPC numbers (Fig. 2D, E, F), suggesting the negative effects of H-LY29 on OPCs are not mediated by PI3K/Akt signalling, but involve other mechanisms, such as ERK1/2, which is almost completely ablated by H-LY29 (Fig. S2G). Equivalent pro-oligodendroglial effects of L-LY29 were observed in MYOLs, with a greater than doubling of the number of PLP+ MYOL in the corpus callosum, compared to controls (Fig. 2G, H; Fig. S2A). Increased numbers of MYOLs were mirrored by a striking increase in the extent of MBP immunolabelling (Fig. 2Gii, Hii), quantified by the MBP+ myelin index, a measure of myelination (Fig. 2K), and increased corpus callosum thickness (Fig. 2L). In contrast, both H-LY29 and TCN halved the numbers of DsRed+ MYOLs (Fig. 2I, J; Fig. S2A), and axonal myelination and corpus callosum thickness were markedly decreased (Fig. 2K, L); these quantitative data correspond to 400% more MYOL and MBP immunolabelling in L-LY29 compared to H-LY29 or TCN. In addition, MYOLs appeared normal in L-LY29 compared to controls, but with increased numbers of cells and myelin sheaths (Fig. 2Gii, Hii), whereas MYOLs were atrophied in H-LY29 and TCN and exhibited abnormal appearing oval somata (Fig. 2Iii, Jii). Due to their lower density, individual MYOLs are more clearly distinguished in the cortex, where they are evidently atrophied and support far fewer myelin sheaths in H-LY29 compared to L-LY29 and controls (Fig. 2M, N, O). Immunolabelling for APC and MBP, which are expressed sequentially in differentiating oligodendrocytes, showed L-LY29 significantly increased the density of both APC+/MBP-‘immature’ oligodendrocytes (imOL) and APC+/MBP+ MYOL (Fig. 2P, Q), whereas the main effect of H-LY29 and TCN was to impair the differentiation of APC+/MBP+ MYOL (Fig. 2Q), consistent with evidence that Akt is required at the onset of oligodendrocyte terminal differentiation (*31*). The results fully validate the pharmacogenomic analysis and demonstrate that LY294002 exerts a profound bipartite concentration-dependent effect on oligodendrocyte lineage cells, with high LY294002 having an anti-oligodendroglial action comparable to inhibition of PI3K/Akt signalling by TCN, whereas low LY294002 massively increased the generation of OPCs and promoted differentiation of myelinating oligodendrocytes by unresolved mechanisms.

**Fig. 2:**
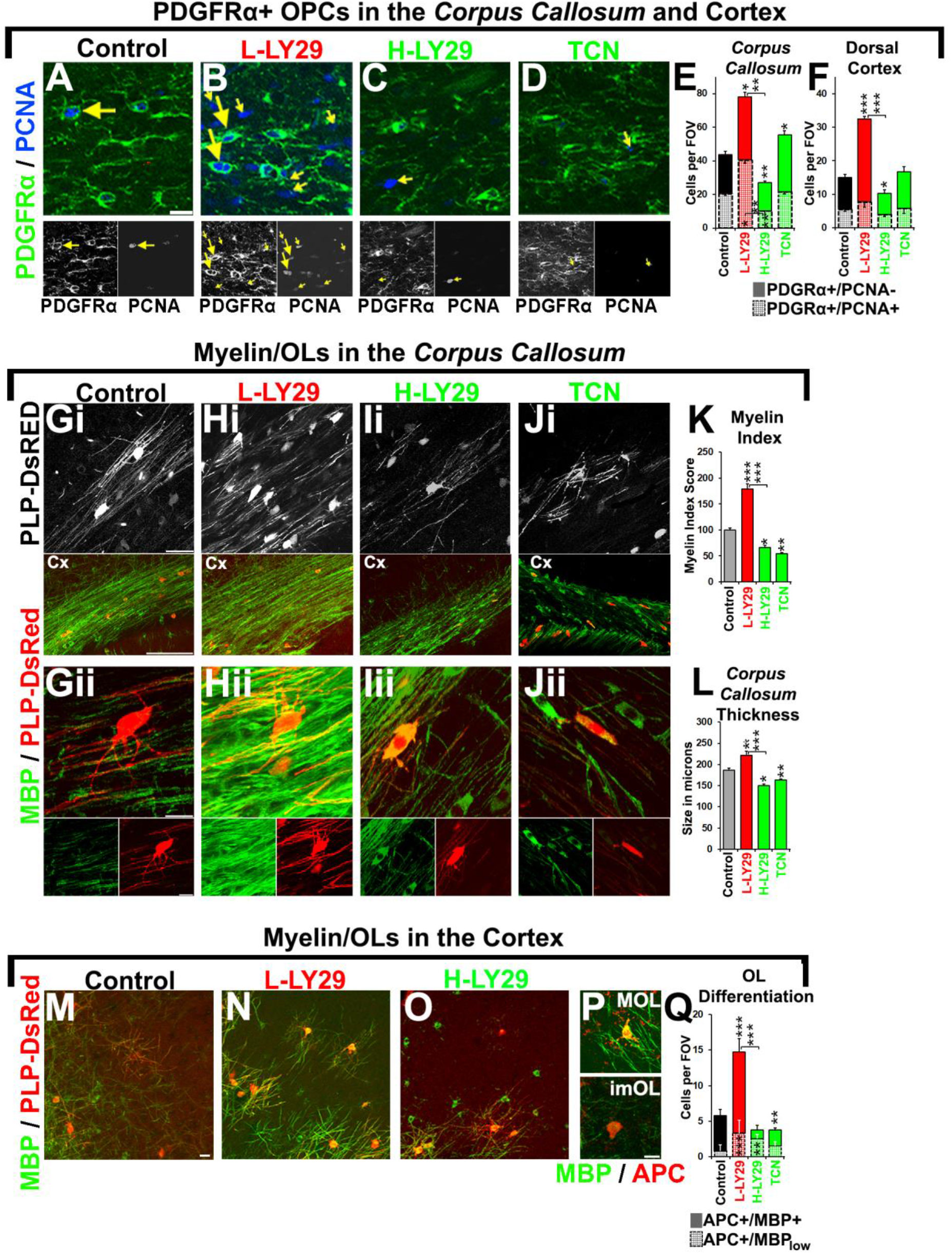
Concentration-dependent effects of LINCS-derived small molecules on oligodendroglia in the periventricular forebrain. Transgenic PLP-DsRed and wildtype P8 mice treated with saline/DMSO as controls, LY-29 and TCN by infusion into the lateral ventricle for 3 days and analysed at P11. Immunostaining done for PDGFRα for OPCs, PCNA for cells in S-phase and MBP for myelin. (**A-F**) Examination/quantification of OPCs in the corpus callosum and cortex and their cell cycle states and quantifications in E. Arrows exemplify PDGFRα+/PCNA+ OPCs and small arrows are PDGFRα-/PCNA+ pre-OPC. Scale bars = 20 μm. (**G-L**) Top panels, middle and lower panels with captions show respectively single z-planes of PLP-DsRed+ myelin sheaths/MYOL, overview of the corpus callosum via MBP immunostainings and higher power cropped confocal sections. Morphologies of OLs induced by L-LY-29 supported further myelin sheaths compared to controls, but OLs appeared abnormal and did not support myelin sheaths in H-LY-29 and TCN. Scale bars = 20 μm in top panels, 150 μm in the corpus callosum overviews and 10 μm in Gii. (**M-Q**) Determination of imOLs and MYOLs densities in the cortex where individual stages of OL units are resolvable as exemplified in P and quantified in Q (n = 4). Flattened confocal z-sections are of 20 μm thickness. Scale bar =20 μm in M and 10 μm in P. In histograms of (E) to (Q), data are mean + SEM quantifications in each region in a constant volume in the case of cell counts (fields of view: FOV); n≥4 animals and each n number represents 3 brain averaged per mouse (***p<0.001, **, p <.01, *, p <.05; t-test). Quantification of myelin index in K or corpus callosum thickness in L are averaged numbers from n = 4 mice (3-4 brain sections per mice) per condition with error bars in SEM. Bonferroni’s posthoc test used to reveal statistically significant differences between the two concentrations of LY-29 and controls.

### LY294002 and TCN regulate oligodendrocyte generation from NSC of the dorsal SVZ

The results above demonstrate that LY294002 regulates the development of forebrain MYOL, which are generated from OPCs that migrate from the dorsal SVZ and are derived from spatially defined pools of NSCs (*2, 32*). We therefore examined the effects of LY294002 on NSC and oligodendroglenesis in the dorsal SVZ, as characterised previously (*32, 33*). NSCs were identified as GFAP+ cells with a radial morphology adjacent to the ventricular surface, either as proliferating NSC (GFAP+/EdU+) or non-proliferating NSC (GFAP+/EdU-) (Fig. 3A, D). Newly generated pre-OPCs/TAPs (transiently amplifying progenitors) were identified by their co-expression of Ascl1 and Olig2 (Olig2+/Ascl1+), whilst OPCs were identified as Olig2+/Ascl1- (Fig. 3B, C, E). Quantification demonstrates that GFAP+/EdU+ and GFAP+/EdU-NSCs were significantly decreased by all three treatments (Fig. 3D), but in addition H-LY29 and TCN treatment severely disrupted NSC morphology (Fig. 3A) and almost completely abolished their proliferation (Fig. 3A, D). In contrast, newly formed oligodendroglial lineage cells exhibited differential responses to treatments compared to controls (Fig. 3Bi, Ci, E), whereby TAPs (Olig2+/Ascl1+) and OPCs (Olig2+/Ascl1-) were markedly increased by L-LY29 (Fig. 3Bii, Cii, E), whereas both populations were significantly reduced by H-LY29 (Fig. 3Biii, Ciii, E), whilst TCN significantly decreased TAPs (Fig. 3Biv, E), but had no significant effect on OPC generation (Fig. 3Civ, E). In addition, quantification of definitive OPC pools in the dorsal SVZ, using immunostaining for PDGFRα and PCNA (as illustrated in Fig. 2A-F), demonstrates their population is more than doubled by L-LY29, whereas they are significantly reduced in H-LY29 and not significantly altered by TCN (Fig. 3F). Overall, these data indicate that low LY294002 drives the generation of oligodendroglial cells from NSCs of the dorsal SVZ and promotes the expansion of newly formed OPC, whereas high hoses of LY294002 and TCN almost completely ablate oligodendrogenesis.

**Fig. 3:**
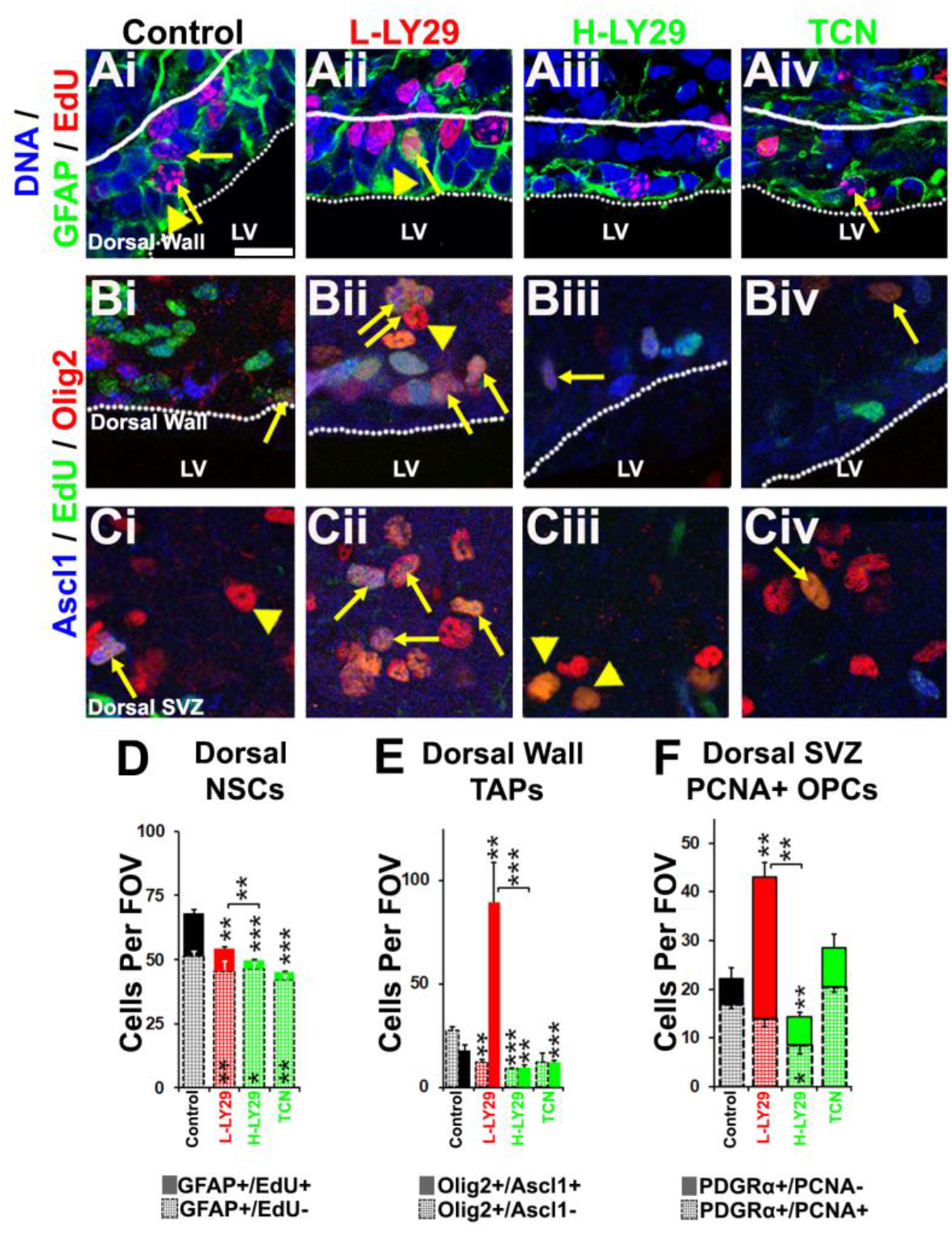
LINCS-derived small molecules differentially modulate oligodendrogenesis. P8 wildtype mice were treated with control (saline/DMSO), LY-29 or TCN by infusion into the lateral ventricle for 3 days and periventricular/SVZ tissue analysed at P11. EdU was given at P9 and P10 by i.p. injections for aiding lineage progression/proliferative cell quantifications in the SVZ. Scale bar = 20 μms. (**A**) GFAP (and staining for EdU for cells that cycled during treatment) for identification of NSCs adjacent to the dorsal ventricular wall. Arrows and arrowheads exemplify NSCs which cycled during treatment and remained quiescent, respectively. Scale bar = 20 microns for micrographs A-C. (**B,C**) Ascl1 and Olig2 for immunolabelling of TAPs and OL lineage cells, respectively, and cells examined directly close to the dorsal wall in B and 70 micron space between the ependymal layer and developing corpus callosum. Arrows and arrowheads exemplify TAPs committed to the oligodendrocytes and OPCs, respectively (Ascl1-/EdU+/Olig2+). (**D to F**) Quantification of dorsal NSCs (D), TAPs (E) and OPCs (E) in the dorsal SVZ tissue and data are mean + SEM in a constant volume (fields of view: FOV); n ≥ 4 animals (***p<0.001, **, p < .01, *, p < .05 t-test/ANOVA where appropriate).

### LY294002 regulates oligodendrocyte generation in developing white matter

Our results demonstrate that LY294002 regulates the generation of oligodendrocytes from the dorsal SVZ. Importantly, endogenous OPCs are a further major source of myelinating oligodendrocytes in the developing and adult brain and are potential targets of small molecules (*21*). To distinguish between the effects of LY294002 on NSCs and endogenous OPCs, we therefore examined the effects of LY294002 *ex vivo* in the optic nerve, a typical white matter tract that contains endogenous OPCs, but not NSCs (*13, 21*), together with the cerebellar slices, using P10-12 SOX10-EGFP reporter mice to identify all oligodendroglial cells (OPC/MYOL), as described previously (*13, 34*). In the optic nerve, LY294002 displayed strict dose-dependent effects, with L-LY29 increasing Sox10+ cells and H-LY29 almost completely ablating oligodendrocytes (Fig. S3A); the pro-oligodendroglial effects of L-LY29 were confirmed in the cerebellar slice, which also enabled qPCR analysis of oligodendroglial transcripts, confirming marked increases in *Sox10* and *Mbp* (Fig. S3B). In the optic nerve and cerebellar slice, OPCs are the sole source of newly generated oligodendrocytes and these results verify that L- and H-LY-29 have a striking bipartite effect on endogenous OPCs, as observed in the forebrain and predicted by pharmacogenomics.

### Transcriptomic profiling of LY294002-responsive signalling pathways that regulate oligodendroglial lineage progression

The adult optic nerve is an excellent model for systems biological analysis of the mechanisms of action of small molecules on glial cells, since it does not contain neuronal nuclei and mRNA transcripts isolated from optic nerves are glial, with insignificant levels from other cells, such as endothelium (*13, 34*). We therefore used a combined neurobiological and transcriptomic analysis of adult mouse optic nerve to determine the effects of LY294002 on oligodendrocyte lineage cells (Fig. 4, SFig. 4). First, we verified that L- and H-LY29 have a bipartite effect on oligodendrocytes in adult white matter, with L-LY29 significantly increasing both Sox10+ OPC/MYOL and PLP+ MYOL, whilst H-LY29 significantly decreased Sox10+ OPC/MYOL, but not PLP+ MYOL (Fig. S4A-D). Notably, H-LY29 markedly inhibited Akt phosphorylation (Fig. S4E), and its negative effect on Sox10+ cells are consistent with PI3K/Akt signalling being essential in OPCs (*31, 35, 36*), whereas the lack of effect of H-LY29 on PLP+ MYOL suggests PI3K/Akt signalling is less important in adult oligodendrocytes. In contrast, the pro-oligodendroglial effects of L-LY29 are at odds with it acting via PI3K/Akt signalling. To resolve this, we performed a differential transcriptomic analysis of optic nerves treated with L-LY29, compared to controls or H-LY29 (Fig. S4F-H). Consistent with the striking bipartite effects of L- and H-LY29, only a relatively small number (85) of genes were common to both treatment groups (Fig. S4H; Tables S1), and STRING and GO analysis highlighted networks and BPs associated with development as being common to L- and H-LY29, with *Igf1* representing a common core hallmark (Fig. S4J). In comparison, differential analysis identified the genes that were regulated by L-LY29 (Fig. S4H), and using the webtool Enrichr the key L-LY29-responsive gene pathways were identified as ‘Focal Adhesion’, ‘Wnt Signaling’ and ‘FAK-PI3K-mTOR signalling’, while BPs induced by L-LY29 included ‘Regulation of cell migration’, ‘Retrograde vesicle-mediated transport’ ‘protein phosphorylation and ‘Mitotic cell cycle phase transition’ (Fig. 4A). To elucidate the oligodendrocyte-specific L-LY29-responsive transcriptional networks, we interrogated our curated expression profiles of OPC- and MYOL-enriched genes (Fig. S4H; see Materials and Methods for details), which are visualised in a NESTED network for exploring the BPs leading to pro-oligodendroglial effects of L-LY29 (Fig. 4B, C). The most significant L-LY29-responsive OPC pathways are associated with cell cycle and differentiation, together with metabolism, nervous system development (p<6.42e-46), Neurogenesis (p<1.07e-37) and Cell morphogenesis (p<1.97e-31). Critical L-LY29-responsive signalling mediators in OPCs are *Egfr* and *Fzd1/2*, with associated key transcriptional regulators *Stat3* and *Tcf4*, indicating key roles of EGFR and Wnt signalling in mediating the pronounced effects of L-LY29 on OPCs, consistent with published evidence (*21, 32, 33, 37, 38*). GO analysis of L-LY29-responsive MYOL genes identified the most prominent biological processes as “Cell Differentiation” (Fig. 4C; Red, *p*<3.68e-08; PPI< 1.54e-12) and “Cellular Protein Metabolic Process” (Fig. 4C; Blue, *p*<2.55e-05), with central roles for *Rhoa* and *Anln* (Anillin), which have established roles in regulating the expression of major myelin proteins (*39, 40*). Network analysis demonstrates that *Rhoa* is at the core of a number of key signalling networks (Fig. 4C; Red, p<3.68e-08; PPI< 1.54e-12), including known pro-oligodendroglial mediators *Fgfr1* (*28*) and *Erbb3* (*27*). The latter regulates oligodendrocytes via RAF-MAPK and PI3K/Akt, both of which were identified by pharmacogenomics as key potential targets for controlling oligodendrogenesis (Fig. S1) and are shown to be directly regulated by LY294002 (Fig. S2). These analyses identify stage-specific signalling pathways by which L-LY29 mediates the observed profound neurobiological effects on oligodendrocytes, with key roles for EGFR and Wnt signalling in massively expanding OPCs and FGFR1 and ERBB signalling in promoting myelination.

**Fig. 4:**
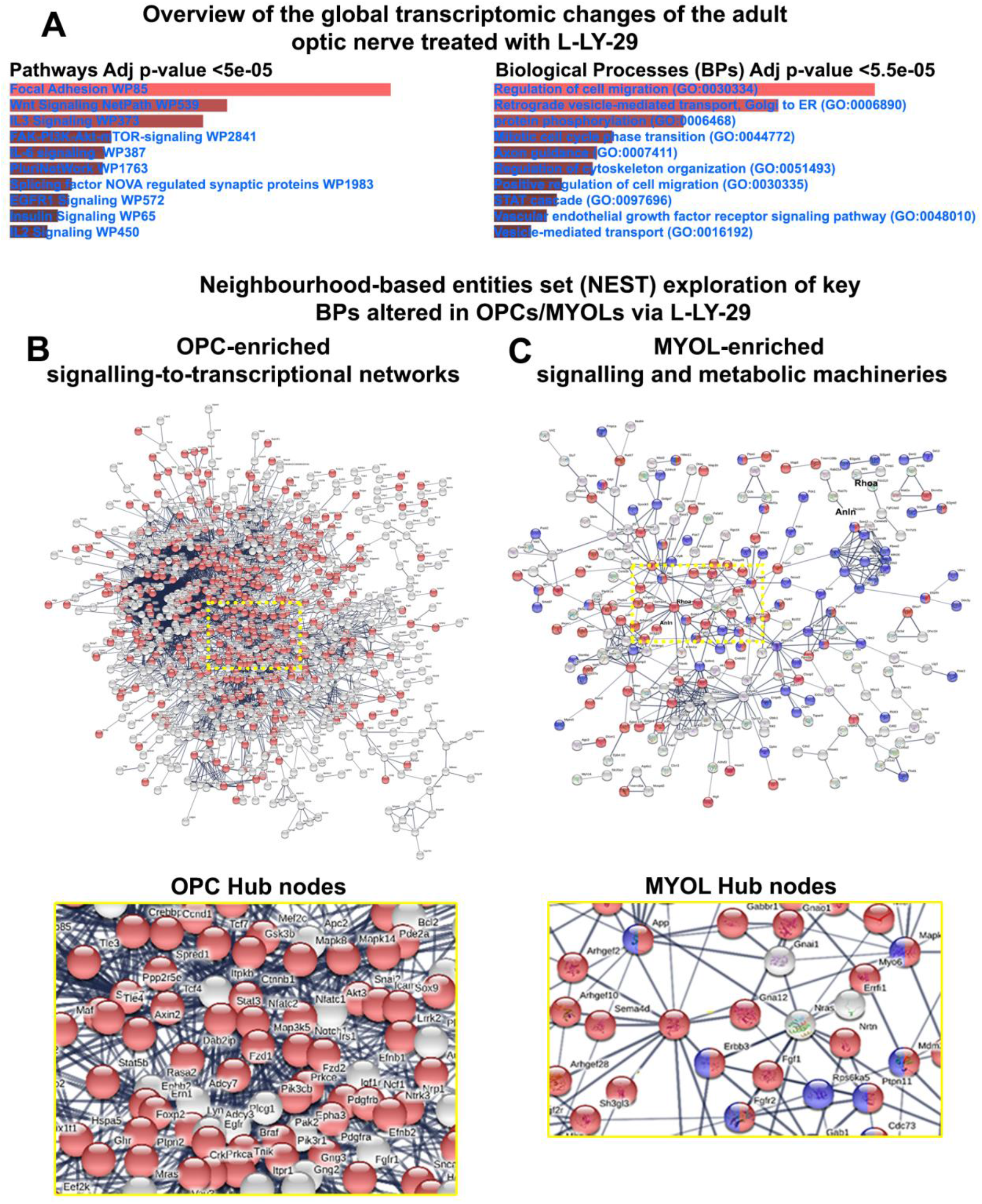
Unravelling of the L-LY29 induced genes in the adult optic nerve and resolving OPC/MYOL cellular networks. (**A**) Significantly differentially expressed transcripts induced by (see also Fig S4H) L-LY29 in the adult optic nerve were inspected for signalling pathways and BP alterations, using the webtool enrichr. The top 10 are presented and ranked by their adjusted (Adj) p-values. (**B,C**) Neighbourhood-based entities set analysis (NEST) of OPC- and MYOL-enriched genes identified by STRING networks for predicted protein–protein interactions (circled in red). Central nodes were cropped and expanded to highlight the essential upstream factors regulated by L-LY29 and nodes are coloured as per key BPs (see corresponding results section for the BPs represented in coloured nodes).

### Validation of systems biology characterization of cell-specific L-LY29-responsive signalling mechanisms

Finally, we performed a validation of the cellular effects predicted by LINCS (SFig. 1) compared to the cellular effects resolved experimentally (Fig. 4, Fig. S4), by interrogating L-LY29-responsive oligodendroglial genes against the LINCS-derived TG pathways and BPs (Fig. 5A). Significantly, LINCS genes enriched in both L-LY29-responsive OPC and MYOL genes were associated with the pathway ‘Focal Adhesion-PI3K-Akt-mTOR-signaling’, which is a key target for pro-oligodendroglial small molecules (Fig. S1). Next, we constructed cell-specific L-LY29-responsive signalling networks in OPCs and MYOLs, using Cytoscape ClueGO (Fig. 5B, C; see Materials and Methods for details). Importantly, the results confirm that pathway terms “Focal Adhesion-PI3K-Akt-mTOR-signaling” and “Focal Adhesion” were both downregulated by L-LY29 in OPCs and MYOLs (Fig. 5B, C). Focal adhesion signal transduction is complex and exerts opposing roles on oligodendrocyte maturation (*41*), whereas signalling networks emanating from PI3K/Akt and affecting signalling by PTEN, GSK3β and mTOR were particularly evident in the effects of L-LY29 on OPCs and are known to be inhibitory in OPCs (*21, 42*). To test the predicted impact of L-LY29 on PI3K/Akt/mTOR signalling, we performed multiplex immunoassays of cerebellar slices treated with L-LY29, which significantly reduced phosphorylation of Akt, mTOR, Pten, p70S6, and Bad (Fig. 5D; *p*<0.05). Thus, these analyses identify stage-specific signalling pathways by which L-LY29 mediates the observed profound neurobiological effects on oligodendrocytes and comprehensively validate the LINCS generated catalogue of small molecules that have the potential to promote oligodendrocyte regeneration and myelination.

**Fig. 5:**
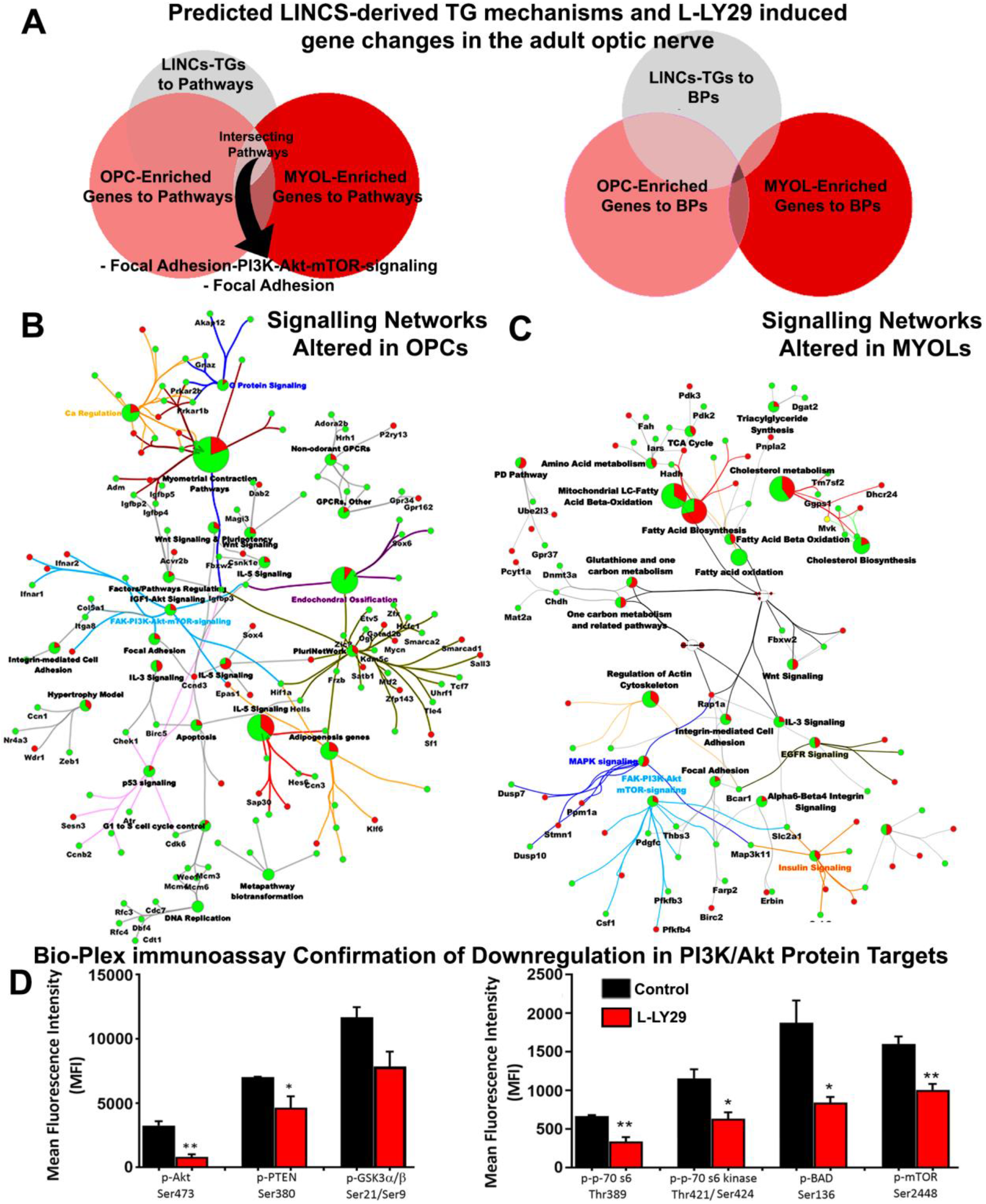
Validation and confirmation of signalling network alterations caused by L-LY29 by phosphoprotein immunoassay. (**A**) Comparison of LINCS-derived TGs processed by Enrichr to identify signalling pathway and BP alterations at two OL lineage stages (OPC-enriched profiles and MYOL-enriched profiles) following L-LY29 treatment of the optic nerve. (**B,C**) Cluego signalling pathway networks of changes in defined stages of OL differentiation. Nodes in red or green represent those that are respectively upregulated or downregulated upon L-LY29. (**D**) Cerebellar slices from P11 mice were maintained in culture for 3 DIV in control medium or medium containing 0.5 μM LY29 and phosphoproteins were assessed by Bio-Plex immunoassay. Data are mean + SEM (n=3 per each group) fluorescence intensity (MFI). phospho-Akt (Ser473), phospho-PTEN (Ser380) and phospho-GSKα/β (Ser21/Ser9); *p<0.05, **p<0.01, two-tailed unpaired t-test.

## Discussion

Connectivity mapping holds considerable potential in the search for new therapies that promote repair in multiple neuropathologies (*3, 13*). However, it is important to determine the potentially contrasting biological actions of drugs depending on the cellular context and their potential dose-dependent effects on intracellular regulatory pathways. Here, using next generation drug connectivity mapping LINCs (https://clue.io) (*11*), we have identified small molecules that have marked dose-dependent effects on oligodendrocytes in the CNS. Surprisingly, LINCS analysis identified the small molecule kinase inhibitor LY294002 as the highest ranking agent predicted to have both pro- and anti-oligodenroglial effects, depending on dose, which we fully validated in multiple neurobiological *in vivo* and *ex vivo* models. Moreover, analysis of LY294002-responsive genes identified the cell-specific mechanisms of action of low and high doses of LY294002 on oligodendrocyte lineage cells. The results demonstrate that a combined drug connectivity mapping and neurobiological strategy is a promising and direct approach to identify small molecules and transcriptional networks that have the potential to promote regeneration and repair in the CNS.

A notable advantage of drug connectivity mapping is the opportunity it provides to identify target gene networks of small molecules and streamline the design of pre-clinical experiments. In the present study, after mapping transcriptional drug responses in iPSC-NSCs onto an oligodendroglial differentiation axis, our meta-analysis of LINCs data revealed that the PI3K/Akt inhibitor LY294002 was predicted to have dose-dependent pro- or anti-oligodendroglial actions. High LY294002 was predicted to cause oligodendrocyte demise via signalling networks and BPs also regulated by the specific Akt inhibitor TCN, e.g. Notch signalling (Fig. S1). As predicted by our connectivity mapping, both H-LY29 and TCN were confirmed to negatively affected oligodendrocyte lineage progression *in vivo* and *ex vivo*, in support of their effects on PI3k/Akt/mTOR signalling (*16, 43, 44*). In contrast, L-LY29 upregulated essential drivers of oligodendrogenesis and differentiation, including Wnt signalling (*32, 33*). These findings were verified *in vivo* in the SVZ and H-LY29 and TCN were shown to perturb common signalling networks in the SVZ that control specification of OPC from NSC, consisting with Akt signalling in dorsal NSC/TAPs directing survival, proliferation, and self-renewal (*45*).

The dorsal SVZ niche during postnatal development expresses over 50 distinct signalling ligands, which are derived from multiple cellular sources within and in the vicinity of the dorsal SVZ, and affect the activity of multiple intracellular pathways, influencing cell fate choices such as survival, self-renewal and commitment to differentiate (*3*). Interestingly, our *in vivo* data reveals complex effects of L-LY29 on multiple stages of oligodendroglial lineage commitment and progression. The observed depletion of proliferating NSCs, accompanied by a dramatic increase in their immediate oligodendrocyte-committed progeny (TAPs), point to a rapid effect of L-LY29 to promote commitment and progression along the oligodendrocyte lineage, consistent with upregulation of cell cycle activity and Wnt signaling (*32, 33*), as predicted by the LINCs TG analysis (Fig. S1A). Importantly, our *ex vivo* data from optic nerve and cerebellar white matter tracts devoid of NSCs and TAPs supported our *in vivo* analyses and demonstrate that L-LY29 promotes OL generation from both the SVZ and endogenous OPCs by triggering a cascade of pro-oligodendrogenic signalling networks, consistent with its broad kinase activity and the pleiotropy of downstream effects of PI3K/Akt/mTOR signalling. LY294002 reversibly inhibits PI3K, but depending on dose can also bind to numerous other proteins (*26*), including the bromodomain proteins BRD2-4 are expressed along the entire oligodendroglial lineage and has the potential to promote oligodendroglial differentiation (*46, 47*). Intriguingly, alterations in GO terms “cell adhesion”, “PTEN signalling”, “Wnt signalling” and “cytoskeletal arrangements” are amongst the most significant transcriptional changes upon partial Brd protein blockade in NSCs (*48*). These data are matched by the major terms predicted and confirmed by our validatory transcriptomic experiments following exposure of optic nerve to L-LY29, lending support to the possibility that targeting Brd proteins could represent a strategy for promoting myelin repair (*49*).

In summary, the extensive datasets hosted by the LINCs consortium, comprising gene expression profiles generated using a diverse set of target cells under a broad set of conditions, provide unparalleled access to the multidimensional interactions emerging when assessing drug-gene interactions at whole-genome levels. Using LINCs, we identified a range of small solutes that have the potential to regulate oligodendrocytes by targeting diverse intracellular signalling pathways. We comprehensively validated our drug networking strategy in multiple *in vivo* and *ex vivo* models, using diverse techniques to analyse the dose-dependent effects of LY294002, including lineage progression characterization, transcriptomics and signalling pathway assays. Our results demonstrated a key role for PI3K/Akt/mTOR signalling in regulating oligodendrocyte generation, as predicted by our analysis of the LINCS dataset, and in support of genetic studies using OPC-specific conditional knockout of mTOR, PTEN and GSK3β (*42*). However, conventional genetic approaches to target specific aspects of intracellular regulatory pathways often yield contradictory findings and cannot resolve dose-dependent effects, which is essential for the development of new therapies (*22, 44, 50*). In contrast, our demonstration of drastically bipartite dose-dependent effects of the PI3K inhibitor LY294002 highlights the power of our pharmacological targeting approach to resolve complex cell-specific signalling networks that is not possible by conventional genetic techniques. These techniques offer unprecedented opportunities to gain insights into hard-to-predict context-specific mechanisms of action of small molecules to promote regeneration and repair in multiple neuropathologies.

## Materials and Methods

### Curation of oligodendroglial hallmark genes from previous studies

Our previous *in silico* screening for therapeutic agents capable of altering developmental myelination was performed by using the first generation of drug connectivity mapping (*3, 51*). The Library of Integrated Network-based Cellular Signatures ((LINCS) (*11*) (https://clue.io), consists of larger and cell-specific resource, comprising 1.3 million expression profiles obtained from over 90 cell lines/IPSC-derived cells, of which the IPSC-NSC drug-induced expression profiles were interrogated. Oligodendrogenesis-associated signatures were compiled using previously generated bulk and single-cell transcriptomic postnatal and adult OL lineage datasets (*17, 18, 52*), together with previously curated ‘pro-oligodendrogenesis’ signatures (*3*). Genes deemed significantly differentially expressed during OL differentiation in these datasets (<5% FDR; >1.8 FC) were standardized into Boolean values while genes commonly expressed within these datasets were removed. The resulting list of 1170 genes comprised the essential landmark genes which define the later stages in the OL lineage as positive values, whereas those in the negative ranges are expressed in dorsal NSCs/TAPs and in the earliest stages of OLs (*3, 13, 17–20*).

### Interrogating the LINCs-database for small molecule acquisition and defining their mechanisms of action on OL lineage cells

The LINCs resource L1000FWD (https://amp.pharm.mssm.edu/l1000fwd/#) was used to process expression signatures to query the IPSC-NPC datasets. The R package g:profiler via RStudio was used to convert mouse gene symbols to human. The following link contains the final expression profiles and derived small molecules: https://amp.pharm.mssm.edu/l1000fwd/result/5e638d9763095f00340d5b7e. A number of small molecules (each tested in triplicate) within the database have been tested under more than one dose/duration condition, thus enabling the dissection of potential concentration-dependent and temporal effects. Duplicates within the top 25 for the pro-oligodendroglial or top 15 anti-correlating (i.e. predicted to inhibit differentiation) small molecules were pooled, averaged and ranked according to the combined pvalues/Z-scores using the R package ggplot2. IPSC-NPC data were downloaded from the L1000FWD resource and tsne coordinates used to construct a geom dot plot via ggplot2, illustrating differences in the transcriptional impact of exposure to the selected small molecules. The R code provided in the resource (https://amp.pharm.mssm.edu/l1000fwd/api_page) was adapted for extracting small molecule target-genes (TGs) in the positive and negative ranges (i.e. increased or decreased upon drug stimulation) among those relevant to the OL lineage/input expression profiles. The TGs for the lower concentration of LY294002 (3.3 μm and 0.37 μm) and the higher concentrations (10 μm, 3 datasets), were merged. TGs for Triciribine were derived from the one available dataset. Next, the TGs were processed for pathway analysis using an R interface of the webtool Enrichr (*53*) (https://cran.r-project.org/web/packages/enrichR/) modified to derive pathways from the extracted small molecule TG lists and visualised using ggplots. The upregulated pathways were maintained in the positive ranges of the combined scoring of pvalues/z-score, whilst for downregulated pathways, values were converted to negative ranges. Pathway terms were shortened to fit within plots and their entire listings, together with raw transcriptomic datasets and output files used in this manuscript will be placed in github (https://github.com/kasumaz) upon acceptance. Files are made available during revision via a cloud drive: https://uni-duesseldorf.sciebo.de/s/7OYQaSbmHTSThy9

### In Vivo Procedures

All animal handling and experimental procedures were conducted in agreement with institutional and regional/national guidelines and the Home Office Animals Scientific Procedures Act (1986) and following approval by the local relevant committees. Animals were housed under standard feeding and lighting conditions. Experiments were performed on the wildtype strain C57/BL6 and on transgenic mouse lines in which fluorescent reporters DsRed or enhanced green fluorescent protein (EGFP) are under control of the oligodendroglial-specific promoters: proteolipid protein 1 (PLP) or Sox10 as characterised previously (*28*). Unless stated, all materials were purchased from Sigma-Aldrich. In vivo experiments were performed on mice aged between postnatal day (P)8 and P11. All procedures were in accordance with the . Mice were either perfused or e killed humanely by cervical dislocation and brains removed rapidly and submerged into ice-cold fixative. Mice aged P8 were treated by intraventricular injections into the lateral ventricle daily for 3 days, and brains sampled at P11 following the final injection. Concentration of injected small molecules into the lateral ventricle were calculated and corrected based on previous spectrophotometry of a GSK3β inhibitor’s bioavailability over time (*21*). Mice were deeply anaesthetised with isofluorane and differing concentrations of LY294002 (Sigma-Aldrich), dissolved in sterile DMSO, sterile-filtered and co-administered with sterile saline delivered into the cerebrospinal fluid (CSF) of the lateral ventricle using a Hamilton syringe, at a point 2 mm from the midline along the Bregma, and to a depth of 2 mm. EdU (5-ethynyl-2’-deoxyuridine) was given as done previously at ages P9 and P10 (32) (*32*).

### Immunohistochemistry

Brains were immersion fixed in 4% paraformaldehyde (PFA) in phosphate buffered saline (PBS), either for 3h at room temperature (RT), or overnight at 4°C. Following fixation, brains were washed in PBS and 50 μm thick coronal sections were serially collected using a vibratome (*28*). Following washes in PBS, a blocking and permeabilization was performed by incubation for 2h at RT or overnight at 4°C in 10% normal goat serum (NGS; Biosera) in 0.3% triton-X-100 in phosphate buffered saline (PBST). Sections were then incubated for 3 h at RT with agitation, or overnight at 4°C, in primary antibodies diluted in NGS: rabbit anti-PDGFRα (1:400, gift from Prof Stallcup); goat anti-PDGFRα (1:200, R&D Systems; mouse anti-APC (CC1; 1:300, Millipore); rabbit anti-GFAP (1:300, DAKO); mouse anti-PCNA (1:400, Sigma-Aldrich) mouse anti-Ascl1 (1:200, BD Biosciences); rabbit anti-Olig2 (1:400, Millipore). After washes in PBST, sections were incubated for 2h at RT or overnight at 4°C in the dark with the appropriate secondary antibodies conjugated with Alexafluor 488, 568 or 405 (1:500, Molecular Probes). Primary antibodies of different origin were diluted together in blocking buffer and co-dilutions of the appropriate secondary antibodies were used. Control experiments were performed using appropriate blocking peptides where available or otherwise by omission of the primary antibody. For PCNA, antigen retrieval was performed by pre-treating sections with PBST and NP-40 1% for 20 min to permeabilize the sections, and following brief washes in PBS, sections were immersed in pre-boiled citric acid and heated in a commercial microwave pressure cooker at full power for 30 sec for 2 cycles. After final washes in PBS, tissues were mounted on poly-lysine-coated glass slides with Vectashield mounting medium (Vector Laboratories) and sealed with coverslips. Images were acquired using an LSM 5 Pascal Axioskop2 or LSM 710 meta confocal microscope (Zeiss). Fluorescence was visualized at 488 nm (green), 568 nm (red) and 405 nm (blue) using argon, HeNe1 and diode lasers respectively, using an x40 oil immersion lens with high numerical aperture (1.3 nm). Optic nerves were immersion fixed in 4% paraformaldehyde (PFA) in phosphate-buffered saline (PBS) for 1 h at RT and following washes in PBS were whole-mounted on microscope slides in VectaShield (VectorLabs).

### Optic nerve tissue and organotypic cerebellar slice cultures

Organotypic cultures of mouse optic nerves were performed as described previously (*13*) and cerebellar slice cultures were prepared using tissue isolated from mice aged postnatal day P10-12 as previously described (*54*). Optic nerves were removed with the retina intact and cerebellar slices were placed immediately in ice-chilled oxygenated artificial (a)CSF composed of: NaCl 133 mM, KCl 3 mM, CaCl2 1.5 mM, NaH2PO4 1.2 mM, HEPES buffer 10mM pH 7.3, 0.5% penicillin and streptomycin (Invitrogen). For optic nerves, n = 6 optic nerves from 3 mice were used per experimental group for confocal microscopy analysis, and 12 nerves from 6 mice were used for transcriptomic analysis, according to power calculations ensuring sample sizes were adequate to detect statistical differences. For preparing cerebellar slices, brain was rapidly removed and placed in oxygenated ice cold slicing solution containing (in mM) 25.95 NaHCO3, 1.39 NaH2PO4, 10 glucose, 124 NaCl, 2.95 KCl, 10 MgCl2, 2 CaCl2, 1 MgSO4, 1000 units/mL penicillin/streptomycin) and 300 μm parasagittal slices were cut using a vibrating microtome 5100 mz (Campden Instruments LTD). Slices were then analysed under the dissecting microscope to ensure maintenance of normal cytoarchitecture. Isolated tissues were carefully cleaned of the arachnoid membrane and any attached peripheral/CNS tissue, then washed in aCSF and placed on semiporous culture membrane inserts (Millipore 0.4 μm). The medium (1 ml) for maintaining optic nerves consisted of 25% horse serum, 49% OptiMEM, 25% Hanks’s balanced salt solution, 0.5% 25 mM glucose, 0.5% penicillin and streptomycin and for cerebellar slices comprised 50% MEM (Eagle) with Glutamax-1, 25% EBSS, 25% horse serum, 130 mM glucose and 1% penicillin-streptomycin (all from GIBCO/Invitrogen). Tissue were maintained ex vivo at 37 °C in 95%O2/5% CO2 for 3 days. LY29 was added directly to the culture medium using the concentrations and duration as stated in the main text and vehicle DMSO used as control. After 3 days for the optic nerve or 7 days for cerebellar slices, tissues were prepared for confocal imaging, RNA extraction or western blot.

### Cell counts, confocal microscopy and image analysis

All experiments were conducted in triplicates and no samples were excluded; due to the study design animals were not blindly selected for group allocation, but all outcome measurements were subsequently conducted blindly, and all samples were included. Periventricular sections containing the lateral ventricle were analysed (>3 sections per brain) using homogenous quantification procedures (*55*); counts of OLs and OPC numbers in untreated controls confirmed that there were no significant differences between sections taken in this area (*21*). Images were captured using a Zeiss LSM Meta 5.1 or Zeiss LSM 7.1 meta confocal microscope and processed with the latest Zeiss ZEN software (black edition), maintaining the acquisition parameters constant to allow comparison between samples. Coronal brain sections were used throughout and cell counts performed in the dorsal SVZ, corpus callosum and cerebral cortex on orthogonally projected confocal z-stacks, of 230 μm2 x 230 μm2 in the x-y-plane, and 30 μm in the z-plane. 1 Hemisphere was used for quantification and the other hemisphere for capturing representative images. For extracellular markers, a nuclear counterstain (DAPI (Invitrogen) or Propidium Iodide (Sigma-Aldrich) was applied to aid quantification.

For analysis of optic nerve cultures, cells expressing either the Sox10-EGFP or PLP-DsRed reporter were visualised at 488nm or 546nm respectively using an argon laser. Images were captured on a Zeiss LSM 710 meta-confocal microscope using a x20 Plan-NEOFLUAR 20 objective with a numerical aperture of 0.50. Images were captured maintaining the acquisition parameters constant between samples. Each nerve counted as a single sample and the total number of cells was counted midway along the length of the optic nerve in a single field of view (FOV), comprising a constant volume of 200 μm × 200 μm in the x–y-plane and 25 μm in the z-plane, commencing 15 μm below the pial surface. For all comparisons, the significance level was set to 5%; due to the explorative nature of this study, no adjustment was made to the significance level. Cell counts are expressed as mean number of cells per FOV ± standard error of the mean (SEM). There were six nerves from three mice in each experimental group and statistical analysis was performed as follows. Measurement of a myelin index was performed on sections from PLP-DsRed mice (21), providing a reliable readout of myelinated sheaths through a z-plane. Confocal micrographs captured with an x40 objective from every 5h confocal section in a series of 30 (n=7 sections of 30 μm thickness) were analysed. The myelin index score presented is the total number of DsRed+ myelin sheaths in a 30 μm thickness from an individual nerve. GraphPad Prism v6 for multiple variables, using either Dunnett’s Multiple Comparisons test, or one-way analysis of variance (ANOVA) followed by Bonferroni’s posthoc test, and for two variable using unpaired t-tests (referred to as t-test) was applied.

### Western blot

P9 pups were bilaterally intraventricularly injected with LY29 as described above. 45 mins following injection, pups were sacrificed by cervical dislocation and tissue rapidly microdissected and flash frozen in lysis buffer in liquid nitrogen for storage at −80°C. Tissue from several pups was pooled to yield individual samples used for later molecular assays. Corpus callosum or optic nerve tissue were centrifuged at 4000 x g to obtain tissue pellets and proteins extracted with lysis buffer as described previously (*21*). After centrifugation for 15 min at 10,000 × g and 4°C, supernatant was transferred to Ultrafree MC centrifugal spin columns (Millipore) for separation and concentration of protein extracts above 15 KDa and Bradford protein assay applied for determination of protein content. For protein analysis, samples were then solubilised and denatured in Lamelli sample buffer (Biorad) with β-mercaptoethanol (Sigma-Aldrich) for 5 min at 95 °C and were placed on ice until loading. 15 μg were loaded onto the gel with Lamelli sample buffer. Solubilised, denatured proteins were then separated via SDS-PAGE and transferred to a PVDF membrane (GE Healthcare, Amersham). Blots were preincubated in a blocking solution of 5% BSA in 0.2% TBST (0.1 M Tris base, 0.1% Tween 20, pH 7.4) for 1 hr at RT, incubated with primary antibodies overnight at 4°C and after washing, with a horseradish peroxidase-conjugated anti-rabbit antibody (1:10,000-1:25,000; Pierce Biotechnology). Primary antibodies were all obtained from Cell Signaling and used in concentrations of 1:500 for phosphor-forms and 1:2000 for total forms of protein. Protein bands were detected by adding SuperSignal West Pico Chemiluminescent Substrate (Pierce) by exposing the blot in a Stella detector (Raytest). Densitometry analysis was performed with NIH software and by normalizing the band intensities to total Akt or total Erk1/2 values and significance assessed using one-way analysis of variance (ANOVA) followed by Bonferroni’s posthoc test.

### RNA extraction and qPCR

RNA extraction from cerebellar slices was performed by placing them in 500 μL of ice-chilled Trizol. 180 μg of total RNA from each sample were converted to single stranded cDNA using the SuperScript^®^ VILOTM cDNA Synthesis Kit (Invitrogen-Life Technologies) following manufacturer instructions. Subsequently, cDNA was added to a mixture of FastStart Essential DNA Probes Master and of FAM dye-labelled primers following manufacturer’s instructions. Samples were run using a Roche Lyghtcycler 96 (Roche) instrument. Reaction consisted of pre-incubation at 95°C for 600 seconds followed by 45 cycles of two step amplification of 95 °C for 10 seconds and 60°C for 30 seconds. Data normalisation to the housekeeping gene Gapdh. Primer sequences: Mbp: 5’ATTCACCGAGGAGAGGCTGGAA’3 / 3’TGTGTGCTTGGAGTCTGTCACC’5; Apc: 5’GTGGACTGTGAGATGTATGGGC’3 / 3’CACAAGTGCTCTCATGCAGCCT’5; Sox10: 5’AGTACCCGCACCTGCACA’3’; / 3’GAAGGGGCGCTTGTCACT’5; Cspg4/NG2: 5’GAGGTCTTGGTGAACTTCACCC’3 / 3’GACAGTAGGAGACCGATGGTGT’5; Gapdh: 5’TTGATGGCAACAATCTCCAC’3 / 3’CGTCCCGTAGACAAAATGGT’5. Relative gene expression levels were determined using the 2 ΔΔ-CT method (*33*). Primers were designed by Primer Express 1.5 software and synthesized by Eurofins MG. Gene expression data are presented as mean and the standard error of the mean (+SEM), and samples were compared for significance via t-test using GraphPad Prism v3.02 software.

### Whole genome transcriptome analysis

Complete details are provided in a recent study from the authors as identical procedures were followed for preparation of RNA for Affymetrix GeneChip Mouse Genome 430 (*13*). Downstream quality control steps and data analysis of produced .CEL image and .CHP image files were performed using Affymetrix GeneChip Operating Software. Agilent GeneSpring GX 12 software was used to normalise the datasets using the MAS-5 algorithm and further statistical analyses. GeneSpring was used to generate hierarchical clustering and the meta-analysis profiles of oligodendroglia-specific (OPC and MYOL) signatures from published databases (*17, 56*). Gene Ontology analysis was performed using ConsensusPathDB, String V10.5 and STITCH db, described in detail previously (*13*). Data are available via the link provided (https://uni-duesseldorf.sciebo.de/s/32r6pWUwjpVG5Z7) until acceptance and raw data are made available in the Github repository.

### Bioplex immunoassay

Cerebellar slices of 300 μm thickness were isolated from P11 wild type mice C57BL/6 strain and incubated for 3d as previously described. After 3d, slices were washed in ice-cold cell wash buffer and lysed according to the manufacturer’s instructions. Total protein content was determined via BCA assay and samples and cell lysate controls were diluted to 10 μg/well (50 μL) in lysis buffer to ensure a constant sample input across all samples. Briefly, samples blank (detection antibody) were loaded in duplicates in a 96-flat bottom well plate (Biorad) with previously diluted custom-made premixed fluorescent beads, which allowed the simultaneous quantification of phosphorylation levels in 7 analytes, and incubated for 15-18h at RT under constant shaking. After washings in wash buffer, samples were incubated in detection antibody for 30 minutes, washed again prior to incubation in Streptavidin-PE for 10minutes. All incubations were performed at RT, under constant shaking and covered from light to protect the light-sensitive beads. Resuspended beads were then transferred to the Bio-plex Suspension Array system (BioRad) for quantification of phosphoproteins fluorescence intensity and results were expressed as mean fluorescence intensity (MFI). Further analysis was performed manually for excluding subtract blank MFI values (background) for each analyte from each sample. Statistical significance was calculated with GraphPad Prism 6.

## Supporting information

All Files

## Supplementary Materials

**Fig. S1:** Prediction of the cellular effects of LINCS-derived small molecules on oligodendrocyte lineage cells.

**Fig. S2:** Concentration-dependent effects of LY294002 (LY-29) and Triciribine on stage-specific oligodendroglia in the corpus callosum and cortex.

**Fig. S3:** LY294002 regulates oligodendroglial cells in the postnatal optic nerve and cerebellar slices ex vivo.

**Fig. S4:** Concentration-dependent effects of LY29 on adult optic nerve oligodendroglia and whole genome profiling for revealing genes regulated by LY29.

**Tables S1:** Concentration-dependency of LY294002 on transcriptome alterations in adult optic nerve. Related to Fig. S4L. Tables can be visualised at the cloud link: https://uni-duesseldorf.sciebo.de/s/L27W5LaxL515JdT

**Tables S2:** Meta-analysis of L-LY294002 on OPC and MYOL enriched expression signatures. Related to Fig. S4L. Tables can be visualised at the cloud link: https://uni-duesseldorf.sciebo.de/s/oaCiiS0VVheergd

## Acknowledgements

We wish to thank the team of Avi Ma’ayan at the Icahn School of Medicine for troubleshooting and repair of the LINCs FWD webtool.

## Authors’ contributions

FP was responsible for the conceptualization, data curation, formal analysis, methodology, supervision. AR contributed to writing, validation, data curation and data analysis. GW carried out data curation, analysis, investigation validation, and methodology. FC was responsible for writing and data analysis. AB was responsible for funding acquisition, project administration, writing, methodology, supervision and validation. KA for funding acquisition, investigation, data curation, methodology, project administration, supervision, validation and writing.

## Data and materials availability

Scripts developed for the first time, Cytoscape files, bulk transcriptomic datasets, gene lists and raw data’s of this study will be placed in Github upon acceptance and a temporary link provided in the methods section.

