## Supplementary material for "Drug connectivity mapping and functional analysis reveals therapeutic small molecules that differentially modulate myelination": All Files

**Fig. S1: Prediction of the cellular effects of LINCS-derived small molecules on oligodendrocyte lineage cells.**

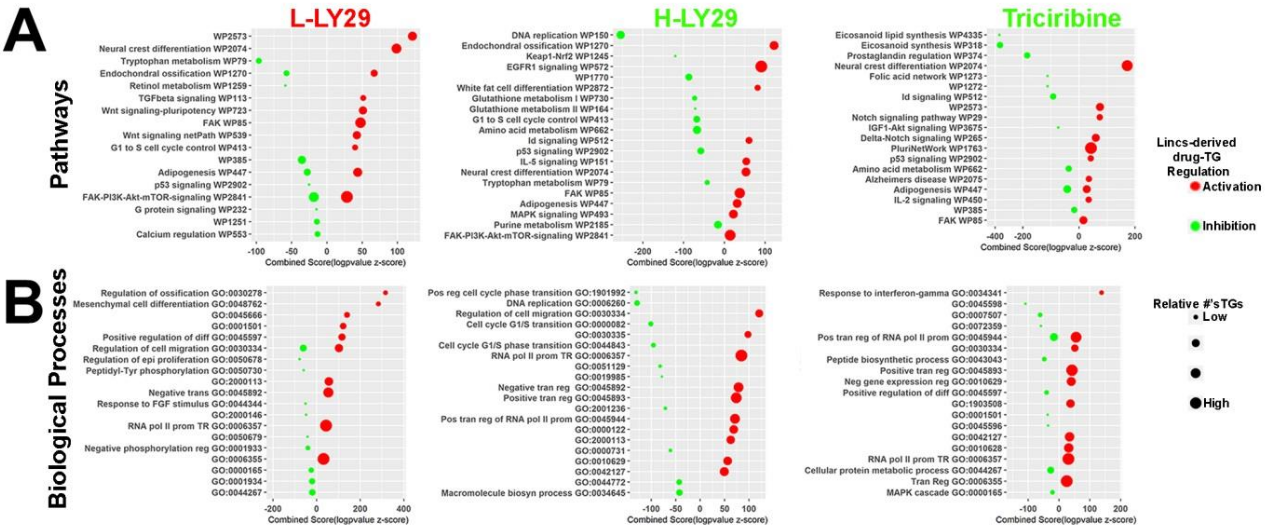


Small molecule TGs Pathway and BP term memberships as revealed by querying (**A**) Wikipathways and (**B**) GO BPs. Geom dot plots illustrating Pathway/BP terms were shortened to fit, arranged by their combined scorings (logpvalue/z-score) and point sizes reflect the relative enrichment. Full list of small molecule TGs, pathways and BPs are available with the raw data.

**Fig. S2: Concentration-dependent effects of LY294002 (LY-29) and Triciribine on stage-specific oligodendroglia in the corpus callosum and cortex.**

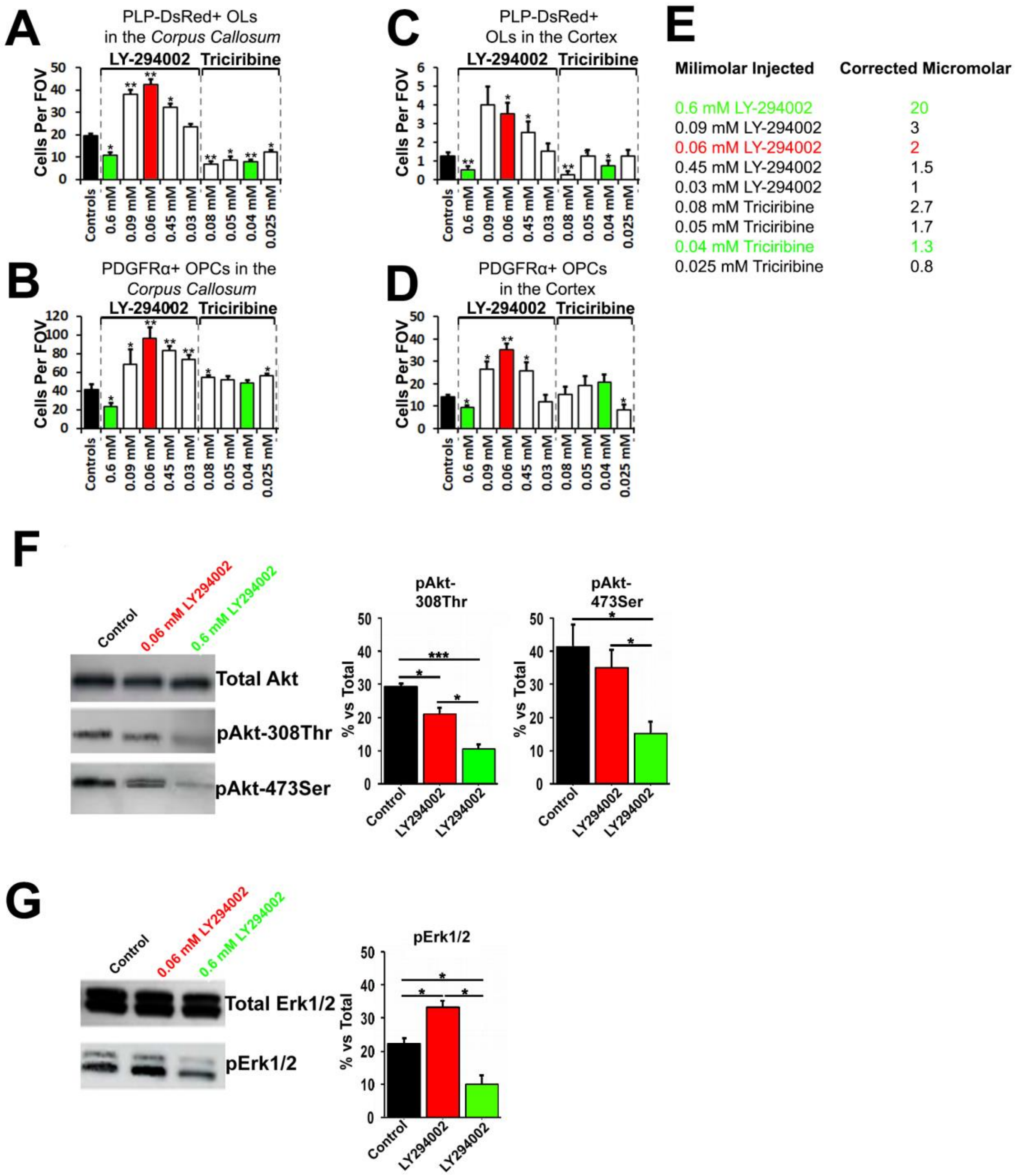


Small molecules were infused at a range of concentrations into the lateral ventricle for 3 days commencing at P8 and the periventricular corpus callosum and cortex were examined at P11. (**A-E**) Abundance of PLP-DsRed-positive oligodendrocytes (**A, C**) and PDGFRα-positive OPCs in wildtype mice (**B,D**) in the specified periventricular tissues. Concentrations are given in **E** as injected and dilution-corrected, based on the measured 20-fold dilution in the CSF (see Methods and Materials). Values are expressed as a mean number of cells per

FOV and error bars represent the SEM (n ≥ 4). *P<0.05, **P<0.01 Dunnett's Multiple Comparisons test. (**F,G**) P9 wildtype mice were treated with 0.06 mM or 0.6 mM LY294002 and saline/DMSO. ~45 min following final infusion, the corpus callosum was rapidly microdissected, snap frozen and pooled in lysis buffer for individual n numbers. (F) Representative immunoblots of total pan-Akt and pAkt at differing sites (~56 KDa) and (G) total Erk1/2 and pErk1/2 (42 and 44 KDa) are shown in controls and different concentrations of LY294002. Signals were analyzed by densitometry versus total protein levels and are expressed in arbitrary absorbance units as the mean ± SEM of 3 independent experiments performed in triplicate from pooled mice. Significance was tested by ANOVA followed by Bonferroni's posthoc test. *p<0.05; ***p<0.001.

**Fig. S3: LY294002 regulates oligodendroglial cells in the postnatal optic nerve and cerebellar slices ex vivo.**

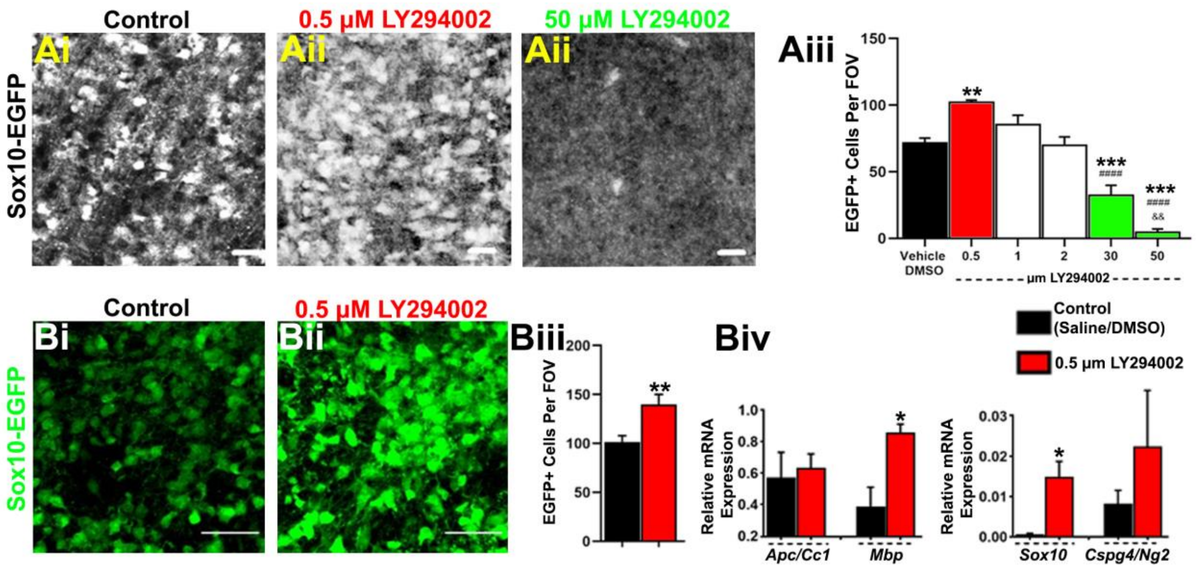


(**Ai-Aiii**) Representative confocal images of optic nerves from postnatal Sox10-EGFP mice maintained in culture for 3DIV, in control medium(vehicle DMSO), and medium containing increasing concentrations of LY-29. Confocal micrographs of the whole-mounted optic nerve illustrates concentration- dependent effects of LY-29 in the adult optic nerve. (**Aiv**) Histogram of mean (+SEM) cell counts per constant FOV (n = 3 for each group); **p<0.01, ***p<0.001 (comparing controls to treatment groups), ####p<0.0001, (comparing LY-29 0.5 µM to higher LY-29 concentrations), One-way ANOVA followed by Dunnett's multiple comparisons test. (**Bi,Bii**) Representative confocal images of cerebellar slices from postnatal Sox10-EGFP mice maintained in culture for 7 d in 0.5 µM LY-29-containing medium; scale bars = 100 microns. (**Biii**) Histogram of mean (+SEM) cell counts per constant FOV (n = 5 for each group). **p<0.01, two-tailed unpaired t-test. (**Biv**) Ex vivo cerebellar slices were maintained in 0.5 µm LY-29 or control vehicle (saline/DMSO) conditions for 3 days and tissue harvested for processing for qPCR for oligodendroglial transcripts. Data are normalised to the housekeeping gene GAPDH (ΔCt) and expressed as mean 2-ΔCt ± SEM (n = 3 for each treatment group). *p<0.05, two-tailed unpaired t-test.

**Fig. S4: Concentration-dependent effects of LY29 on adult optic nerve oligodendroglia and whole genome profiling for revealing genes regulated by LY29.**

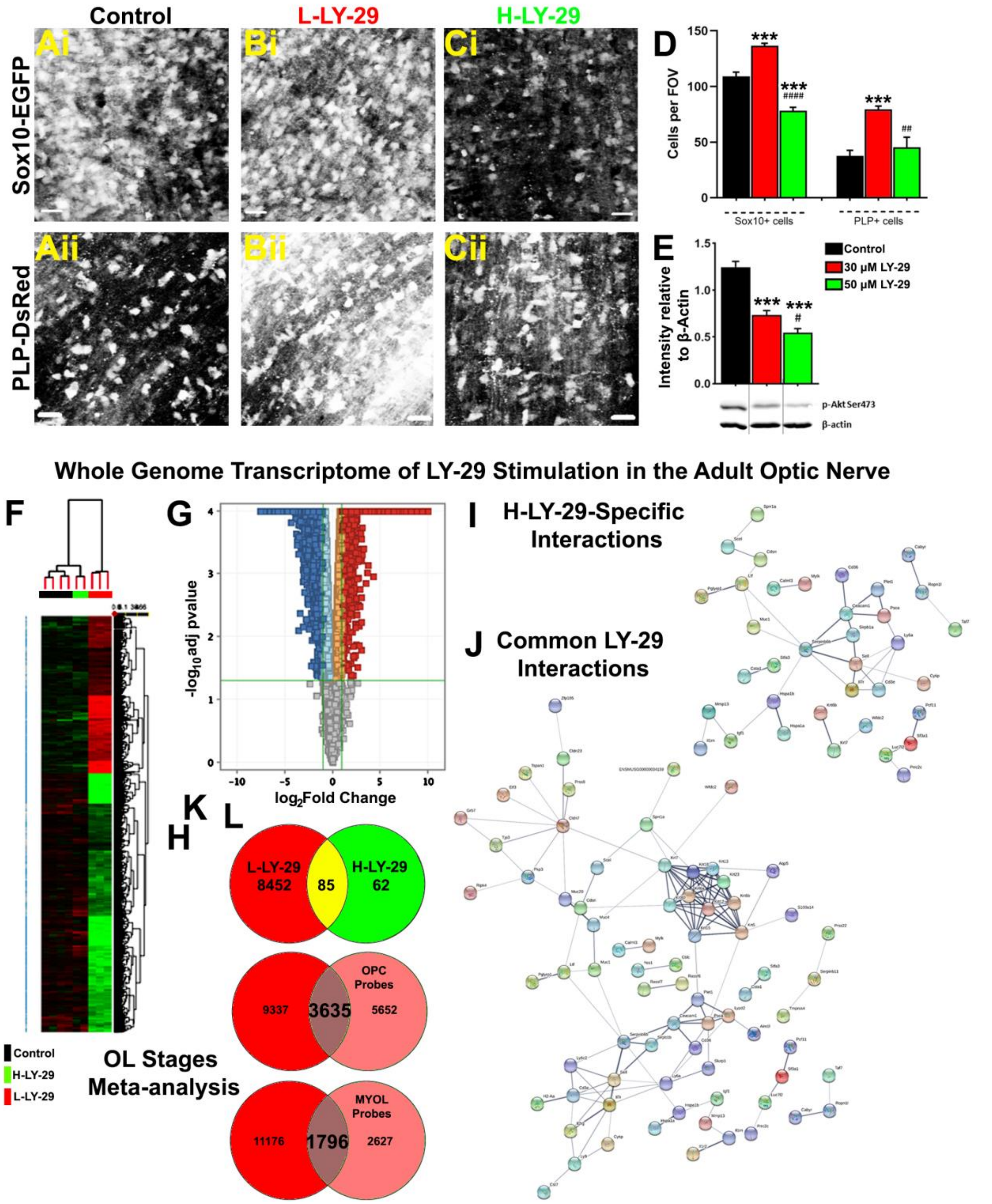


(**A-C**) Representative confocal images from Sox10-EGFP (top panels) and PLP-DsRed1 (lower panels) optic nerves maintained in culture for 3 d in control (vehicle DMSO medium) (A), 30 µM LY29 (B), or 50 µM LY29 (C); scale bars = 20 µm. (**D**) Histogram of mean (+SEM) cell counts per constant FOV (n = 6 for each group), ***p<0.001, (comparing vehicle control to all treatments), ##p<0.01, ####p<0.0001 (comparing 30 µM LY29 to 50 µM), One-way ANOVA followed by Tukey's post-hoc test. (**E**) Relative protein levels of p-Akt-Ser473 in optic nerves incubated for 1 hour either in control aCSF, 30 µM LY29, or 50 µM LY29 were measured by western blot and normalized to β-actin. Quantification of mean ± SEM band intensity relative to β-actin, n = 3 replicates for each data point (n = 6 optic nerves per pooled replicate); ***p<0.001 (comparing control to all treatments) One-way ANOVA followed by Dunnett's post-hoc test, #p<0.05 (comparing 30 µM LY29 to 50 µM), two-tailed unpaired t test. (**F-G**) Heatmap of transcriptional changes occurring in the adult optic nerves incubated in control medium and in 30 or 50 µM LY29. Red, green and black represent probes upregulated, downregulated or showing no change, respectively, and the most significant probes shown on a volcano plot in G. (**H**) Venn diagram of probes specific to or common between the concentrations of LY-29 (top Venn's), meta-analysis of OPC-enriched (middle Venn's) or MYOL-enriched (bottom Venn's) profiles using publically available OPC- and MYOL-derived datasets (see Materials and Methods). (**I-J**) Protein interactome networks generated using the STRING database of genes common to both LY29 concentrations or those specific to H-LY29.
